# Biochemical profile and bioactive potential of wild folk medicinal plants of Zygophyllaceae from Balochistan, Pakistan

**DOI:** 10.1101/2020.03.30.016212

**Authors:** Alia Ahmed, Amjad Hameed, Shazia Saeed

## Abstract

Recent focus is on analysis of biological activities of extracts from plant species. Zygophyllaceae is exceedingly important angiosperm family with many taxa being used in folk medicines widely dispersed in arid and semi-arid zones of Balochistan, Pakistan. Only a small proportion of them have been scientifically analyzed and many species are nearly facing extinction. Therefore present investigation explores the biochemical and bioactive potential of fourteen folk medicinal plants usually used for treatments of different ailments. Fresh aerial parts of nine taxa and two fruit samples were collected from plants growing in arid and semi-arid zones of Balochistan and analyzed for enzymatic, non-enzymatic and other biochemical activities. Higher phytochemical activities were detected in the aerial parts. Superoxide dismutase was detected maximum in *Fagonia indica*, (184.7±5.17 units/g), ascorbate peroxidase in *Tribulus longipetalus* subsp. *longipetalus* (947.5±12.5 Units/g), catalase and peroxidase was higher in *Peganum harmala* (555.0±5.0 and 2597.8±0.4 units/g respectively). Maximum esterase and alpha amylase activity was found in *Zygophyllum fabago* (14.3±0.44 and 140±18.8 mg/g respectively). Flavonoid content was high in *T. longipetalus* subsp. *longipetalus* (666.1±49 μg/ml). The highest total phenolic content and tannin was revealed in *F. olivieri* (72125±425 and 37050±1900 μM/g. respectively). Highest value of ascorbic acid was depicted in *F. bruguieri* var. *rechingeri* (448±1.5 μg/g). Total soluble Proteins and reducing sugars were detected higher in *P. harmala* (372.3±54 and 5.9±0.1 mg/g respectively). Maximum total antioxidant capacity (TAC) was depicted in Z*. simplex* (16.9±0.01 μM/g). Pigment analysis exhibited the high value of lycopene and total carotenoids in *T. terrestris* (7.44±0.2 and 35.5±0.0 mg/g respectively). Chlorophyll a, b and total chlorophyll content was found maximum in *T. longipetalus* subsp. *pterophorus* (549.1±9.9, 154.3±10 and 703.4±20.2 ug/g respectively). All taxa exhibited anti-inflammatory activity as well as anti-diabetic inhibitory potential. Seed extracts of *Zygophyllum eurypterum* (96%) exhibited highest inhibitory potential, along with twelve other taxa of Zygophyllaceae indicated (96-76%) activity when compared with the standard drug diclofenac sodium (79%). Seeds of *T. longipetalus* subsp. *longipetalus* (85%) exhibited the highest anti-diabetic activity; other eleven taxa also exhibited inhibitory activity of α-amylase ranging from (85-69%) compared with Metformin (67%) standard drug. Phytochemical screening revealed that selected taxa proved to be the potential source of natural antioxidants and could further be explored for in-vivo studies and utilized in pharmaceutical industries as potent therapeutic agents validating their ethno-pharmacological uses.

## Introduction

Most of the world’s population, particularly in the underdeveloped and developing countries, depends mainly on herbal medicines to cure various ailments [1]. Wild medicinal plants are being utilized as folk medicines globally. Numerous plant parts have various therapeutic properties and utilized as herbal medicines to treat various disorders and infections. Currently most prevailing and powerful drugs used were derived from medicinal plants [2,3] Thus, for biological and antioxidant capacity plant based derivatives must be explored. During the last few years herbal medicines and their derivatives seek much attention because of their traditional use. People of many developing countries depended on the medicinal plants for their healthcare. Focus on antioxidants specifically of herbal medicines originates from their ethnopharmacological utilization [4]. Many medicinal plants extract constituted various types of chemicals, each have a property to control a variety of biological and pharmacological activities such as antimicrobial, anti-parasitic, anti-diabetic, antioxidant, anti-inflammatory and anticholinesterase. These chemicals may have a combined effect in controlling various diseases [5,6]. Diabetes mellitus is commonly related with metabolic disorders linked with numerous macro and micro vascular problems. These disorders accelerate morbidity and mortality [7–10]. It has also been revealed that oxidative stress, rise in free radicals and deterioration of antioxidant defense may mediate the prevalence of diabetes associated complications in diabetic patients [11,12]. Many degenerative disorders like rheumatoid arthritis, joints and shoulder inflammation, heart disease, muscular inflammation, asthma, cancer, and inflammation of gastrointestinal tract commonly related with inflammatory processes [13,14].

During various metabolic processes free radicals formed in the living systems. The reactive oxygen species (ROS) comprised mostly of hydroxyl radical (• OH), superoxide anion (O_2_^−^) and hydrogen peroxide (H_2_O_2_) formed in little quantity for diverse physiological processes [15]. Though, increased amount of these radicals resulted in various disorders like, Alzheimer disease, atherosclerosis, inflammation of various organs and cancer [16]. Oxidative stress, resulting from excessive ROS, causes detrimental effects on biomolecules, leading to many pathological and neurological disorders. The integrated antioxidant systems can balance the toxicity of oxidative reactive species which comprises of enzymatic and non-enzymatic antioxidants. Enzymatic antioxidants such as superoxide dismutase (SOD), peroxidase (POD), ascorbate peroxidase (APX) and catalase (CAT) remove hydrogen peroxide, scavenging free radicals and intermediates of oxygen [17]. Non-enzymatic antioxidants comprised of lipid soluble related with membrane like alpha tocopherol, beta carotene and water soluble reducers like ascorbate, glutathione and phenolic [18]. The antioxidants like polyphenols safeguard the biological systems and oxidative processes. Therefore, controlling diseases induced with oxidative stress by the consumption of herbal products as a rich source of phyto-antioxidants [19]. It has been proved that herbal extracts containing phyto-antioxidants particularly polyphenols, flavonoids, tannins and other associated compounds have progressive health effects and decrease disease risk. Recent researches revealed effects of bioactive and antioxidant potentials of medicinal plants. [20,21].

The Zygophyllaceae consists of diverse habits of wild flora including succulents, herbs, undershrub, shrubs and small trees. The habitat of these plants is predominantly desert or saline areas of temperate and tropical regions around the globe [22,23]. *Tribulus, Fagonia, Zygophyllum* and *Peganum* are being considered as the main genera. Many taxa of the family being used ethnomedicinally. Ref ?

*Fagonia* Linn., is a genus of wild, flowering plants of the family, Zygophyllaceae, having about 45 species all over the world. The distribution of the genus includes parts of Africa, the Mediterranean Basin, Asia and parts of the America. In Pakistan 10 species are reported earlier, in Balochistan it has 7 species present. While working in the field of southern Balochistan we collected 7 species including few sub-species of the genus.

*P. harmala* Linn., is a perennial herbaceous plant usually 25-60 cm tall yellowish white flowers appears in April to May and commonly dispersed in many regions of the province.

Genus *Tribulus* L. is considered as the most complex genus in Zygophyllaceae because of enormous number of invalid specific epithets and also of the variations present in various populations In Pakistan 4 species are reported.

*Zygophyllum* Linn., with about 90 species, grows mainly between northern Africa and central Asia mainly in arid and semiarid areas.

Keeping in view the significance of the medicinal plants of Zygophyllaceae, the present study was conducted and most probably the first comprehensive report on phytochemical screening of leading genera of Zygophyllaceae from the arid and semi-arid regions of Balochistan, Pakistan. The specific objective was to evaluate the bioactive potential and to detect the anti-diabetic and anti-inflammatory activities of aerial parts and seed extracts of these medicinal plants to validate their folk-uses and subsequent utilization as a source of novel drugs.

## Materials and Methods

### Plant collection

The selection of medicinal plants was based on ethno-botanical appraisal. Wild plants were collected from different areas of Balochistan, Pakistan. Voucher specimens were prepared, identified and submitted in Botanical garden Herbarium university of Balochistan, Quetta. Furthermore records also available in open herbarium for future reference (www.openherbarium.org). Fresh aerial parts of nine plants, fruits of two species of *Tribulus,* dry aerial parts of fourteen plants and ten seeds were collected from different ecological zones of Balochistan (Table 1). Fresh plant samples were transported and kept at −20°C. Samples were collected and shade dried and stored at room temperature. Experiments were conducted at Plant Breeding and Genetics Division (MAB Lab) Nuclear Institute for Agriculture and Biology (NIAB), Faisalabad, Pakistan.

Table 1: List of selected taxa of Zygophyllaceae with geographical coordinates of the collection sites and voucher specimen’s number

### Extraction of antioxidant enzymes

Fresh aerial parts of nine plants, two fruits, dry aerial parts and seeds (0.15g) were subjected in 1ml phosphate buffer (50 mM, pH 7.8) to ground. Further the mixture was centrifuged at 14,000×g (20 min, 4°C). Now the supernatant of this extracted plant material used to perform further phytochemical activities. All the data were taken in replicates of three.

### Superoxide dismutase (SOD) assay

Method of [24] used to determine SOD activity by homogenizing the fresh aerial parts and fruits of selected taxa in phosphate buffer (50 mM, pH 7.8), EDTA (0.1 mM) and DTT (1 mM) following the procedure of [24]. Further analyzed by assessing its property to stop the photochemical reduction of nitro-blue tetrazolium as explicated by [25]. One unit of SOD activity was demarcated as the amount of enzyme causing 50% inhibition of photochemical reduction of nitro-blue tetrazolium.

### Peroxidase (POD) assay

The assessment of POD activity carried out using method [26] was followed with few changes. Homogenized mixture of the aerial parts and fruits prepared in1 ml phosphate buffer (50 mM, pH 7.8), EDTA (0.1 mM) and DTT (1 mM). The assay solution contained 535μl distilled water, phosphate buffer 50 mM (pH 7.0), Guaiacol (20 mM), H_2_O_2_ (40 mM) and 15μl enzyme extract. The addition of enzyme extract initiated the reaction. At 470 nm absorbance rises was noted at interval of 20 sec. Absorbance change of 0.01 min^−1^ was demarcated as one unit POD activity. Enzyme activity was expressed on the basis of fresh sample weight.

### Catalase (CAT) assay

Catalase activity was measured by homogenizing the aerial parts and fruit samples prepared in phosphate buffer (50 mM, pH 7.8), EDTA (0.1 mM) and DTT (1 mM). CAT was assessed by following the method of [26]. The activity was measured in a solution contained 59 mM H_2_O_2_ and 0.1 ml enzyme extract. At 240 nm decrease in absorbance recorded after interval of 20 seconds. Change in absorbance of 0.01 per min defined CAT activity of one unit. Enzyme activity was expressed on fresh weight basis.

### Ascorbate peroxidase (APX) assay

The assessment of APX activity carried out by following the [24] method. Samples were extracted in phosphate buffer (50 mM, pH 7.0). APX activity was analyzed by the method of [24]. For the measurement of APX the assay buffer contained potassium phosphate buffer (200 mM, pH 7.0), EDTA (0.5 M), ascorbate (10 mM), 1 ml of H_2_O_2_ and 50μl supernatant. At 290 nm absorbance decrease was noted after every thirty seconds to estimate the oxidation rate of ascorbic acid [27].

### Hydrolytic Enzymes

#### Esterase activity

The α-esterase was determined by using the method [28]. α-naphthyl acetate used as substrates, the reaction mixture contained 30 mM α-naphthyl acetate (30 mM), acetone (1%), and phosphate buffer (0.04 M,pH=7) and enzyme extract. Incubate this mixture for 15 min at 27°C in dark. After 15 min added staining solution 1 ml (Fast blue BB 1% and SDS 5% with ratio of 2:5) and again incubate in dark for (20 min, 27 °C). Absorbance at 590nm was measured for α-naphthol produced amount. Using standard curve, enzyme activity was α-naphthol produced in μM min^−1^/g wt.

#### Alpha amylase activity

A modified method for alpha amylase activity was followed for all plant samples as described by [29].

### Other biochemical parameters

#### Total oxidant status (TOS)

Method of [30] used to determined TOS. The assay established on ferrous ion oxidation into ferric ion. The presence of oxidants in the sample in acidic medium and ferric ion measurement produced by xylenol orange [31]. The assay based on two mixtures R1 (stock xylenol orange solution (0.38g in 500μL of 25Mm H_2_SO_4_) 0.4g NaCl, 500 μL glycerol and volume up to 50Ml with 25mM H_2_SO_4,_ sample extract and R2 (0.0317 g o-dianisidine, 0.0196 g ferrous ammonium sulphate (II). Absorption measured at 560nm after 5 min by using spectrophotometer.

#### Pigment analysis

The concentration of Lycopene, chlorophyll (*a* and *b*), Total chlorophyll and carotenoids were examined by method of [32]. Samples (0.2 g) were grind in acetone (80%) and centrifuged at 10,000 g for 5 minutes. Absorbance measured at 645, 663 and 480 nm by using a spectrophotometer.

#### Total phenolic contents (TPC) and Tannin

Micro colorimetric assay was used to measure TPC, by using Folin-Ciocalteu (F-C) reagent [33] with some modifications.0.05 g sample was kept in 95% methanol in dark for 48 hours. After 48 hours take the supernatant add 150 μl FC reagent (10%) and 1.2 ml sodium carbonate (700 mM). Place this mixture at room temperature for 1 hour and take reading at 765 nm. Linear regression equation was calculated by using standard curve of Gallic acid at different concentrations. To measure Tannin add (0.1g) PVPP and in above prepared sample vortex vigorously and centrifuge again at 14000 rpm, measured reading at 765nm.

#### Determination of Total Flavonoid Content

Assay was determined by colorimetric method using Quercetin as standard. Take 200 μl sample prepared in 95% methanol extract and phosphate buffer (40 mM, pH 6.8). Add 50 μl AlCl_2_ (10%) 50 μl Potassium Acetate (1M) Incubate the mixture at room temperature for 40 min and take reading at 415nm absorbance.

#### Total Antioxidant Capacity

A modified method of TAC was followed as described by [34]. Due to the presence of antioxidants in sample, ABTS assay represents decrease of 2, 2-azino-bis (3-ethylbenzothiazoline-6-sulfonate) radical cation ABTS•+ (blue-green in color) into original ABTS (colorless compound). The antioxidants of the sample extract according to their content decolorize the ABTS•+ radical cation. The reaction mixture contained reagent R1, sample extract and reagent R2. After 5 min at wavelength of 660nm, the absorption of each reaction mixture was measured. This analysis used AsA (ascorbic acid) to develop a calibration curve. The results for antioxidant contents found in plant extracts were measured as μM AsA equivalent to1g.

#### Reducing Sugars (Sugar Content)

Assessment of reducing sugars level in plant samples was determined by dinitrosalicylic acid method proposed by [35].

#### Total soluble protein content

Protein estimation of plant samples was based on quantitative protein analysis described by [36]. Aerial parts and fruits samples were homogenized in potassium phosphate (50 mM, pH 7.0). Supernatant 5μl and NaCl (0.1N) mixed with1.0 ml of Bradford dye. Incubate the mixture for 30 min to get a protein dye complex. Measure the quantity at 595 nm absorbance by spectrophotometer.

### *In Vitro* Anti-Diabetic Activity (enzyme α-amylase inhibition method)

The *in Vitro* anti-diabetic activity was determined by assaying the inhibitory activity of the enzyme α-amylase which involves in the breakdown of starch to produce glucose [37]. In this method, 1 ml of methanolic extracts of all species were tested separately and thus added to 1 ml of the enzyme α-amylase in a test-tube and incubated for 10 min at 37°C. Then 1 ml of 1% starch solution was added into it and again incubated for 15 min at 37°C. Then 2 ml 3, 5-dinitrosalicylic acid reagent was added into it, in order to terminate the reaction. The reaction mixture was then incubated in boiling water bath for 5 min and then allows it to cool at room temperature. The absorbance of the reaction mixture was then measured at 546 nm in a spectrophotometer. The standard (control) of the reaction without the extract represents the 100% enzyme activity. The % age inhibition of enzyme activity of α-amylase was determined by:

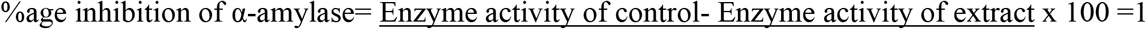

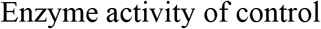

### *In Vitro* Anti-Inflammatory Activity (Protein Denaturation Method)

The protein denaturation assay was determined using a modified method as described by [38]. Briefly, the reaction mixture (0.5 mL; pH 6.3) consisted of 0.45 mL of bovine serum albumin (5% aqueous solution) and 0.05 mL of distilled water. The pH was adjusted to 6.3 using a small amount of 1 N HCL. 1 mL of acetone or aqueous extract with final concentrations of (0.1 to 0.5 mg/mL) was added to the reaction mixture. These were incubated at 37°C for 30 min and then heated at 57°C for 5 min. After cooling the samples, 2.5 mL of phosphate buffer solution (pH 6.4) was added. Turbidity was measured spectrophotometrically at 660 nm. For the negative control, 0.05 mL of distilled water and 0.45 mL of bovine serum albumin were used. Diclofenac sodium with the final concentration of 100, 200, 300, 400, and 500 *μ*g/mL was used as reference drug. The percentage inhibition of protein denaturation was calculated by using the following formula:

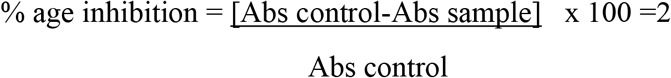

## Statistical Analysis

Data was recorded in mean±SEM. Resulting data were analyzed by applying descriptive statistics. Two-way ANOVA with three replications were used in analyses. Significance of data was tested by analysis of variance and Tukey (HSD) Test at p<0.001using XL-STAT software.

## Results and Discussion

Present study is based on analysis of biological activities of extracts from fifteen plant species of Zygophyllaceae, remarkably important angiosperm family with many taxa being used in folk medicines. There are not much data available in literature about these plants so making a comparison of the results we obtained is not easy. Nonetheless, a few papers reported few biological activities of *T. terrestris*, *P. harmala* and few species of *Fagonia*. Present investigation explores the presence of enzymatic constituents such as SOD, POD, APX, CAT, esterase, alpha amylase, non-enzymatic antioxidants and other phytochemical like AsA, TOS, TAC, TSP, TPC, TF, tannin and pigments. Selected plants also provided evidence for anti-diabetic and anti-inflammatory potential of varying extent in seeds and aerial parts. The difference in pro-oxidants and antioxidants causes oxidative stress and chronic diseases in the body [39]. Cellular damage results in causing cancer. One of the mechanism behind the anti-oxidation is free radical scavenging action [40]. POD helps in scavenging the reactive oxygen species (ROS), causing cell oxidative injury [41]. Highest value of peroxidase and catalase were depicted in the aerial parts of *P. harmala* (2597.8±0.4 units/g f. wt. and 555.0±5.0 Units/g f. wt. respectively) show in S. Fig 1a and b respectively. Plant species having high antioxidant activities can be utilized for different therapeutic applications for treatment of oxidative stress induced diseases.

Antioxidant enzyme catalase present in all animal tissues with its highest activities in liver and red blood cells which defends the tissues from highly reactive hydroxyl radicals by decomposing hydrogen peroxide. Decrease amount of catalase causes numerous damages due to hydrogen peroxide and superoxide radical assimilation [42]. Superoxide dismutase enzyme referred as the significantly involved in cellular defense, therefore it is considered as an indicator of antioxidant capacity [43]. The highest value of superoxide dismutase (184.7±5.17 units/g f. wt.) was observed in *F. indica* shown in S. Fig. 1c. Traditionally this plant being used for anticancer treatment and possibly detected high concentrations of SOD and TAC may be responsible for this therapeutic effect.

Highest APX value was recorded in *T. longipetalus* subsp. *longipetalus* (947.5±12.5 Units/g f. wt.) S. Fig. 1d. Ascorbate peroxidase (APX) enzyme is crucial for the protection from damage by H_2_O_2_ and hydroxyl radicals (•OH) [44]. Antioxidant enzymes activities namely, CAT, POD and SOD were reported earlier in *Rumex obtusifolius* a wild medicinal plants and found to have good antioxidant capacity [45]. Similarly, *Calamintha officinalis* also has potent antioxidants [46]. Alpha amylase and esterase activity was higher in *Z. fabago* (140±18.8 mg/g. and 14.3±0.44 μM/min/g f. wt. respectively) S. Fig. 1e and f respectively. Esterase plays an important role in the disintegration of natural constituents and industrial pollutants and other toxic chemicals. It is also beneficial for the production of optically pure compounds, perfumes, and antioxidants [47].

**S. Fig. 1:** comparison of a) Peroxidase Activity b) Catalase Activity c) Superoxide Dismutase Activity d) Ascorbate Peroxidase Activity e) Alpha Amylase f) Esterase activity.

Maximum TAC was depicted in fresh samples of *Z. simplex* (16.9±0.01μM/g. f. wt.) followed by *F. indica* (15.6±0.04μM/g. f. wt.) shown S. Fig. 2a. No significant variation was detected among all dry aerial parts of selected plants. In general maximum TAC was detected in seed of *Z. eurypterum* (15.8±2.2 μM/g. dry wt.) followed by aerial parts of *Z. simplex* (15.7±2.33 μM/g.dry wt.). In seeds of selected taxa no significant variation was detected the maximum TAC was in *F. olivieri* (15.6±2.4 μM/g. s. wt.) second highest was in *F. bruguieri* var. *rechingeri* (15.5±2.5 μM/g. s. wt.) Previously aerial parts of *F. longispina* reported to be a good source for natural antioxidants [48]. Previously *F. cretica* was also found to have high antioxidant and radical scavenging potential due to high TPC and TFC [49]. *F. olivieri* can serve as a natural source to develop the free radical scavengers beneficial in the prevention of oxidative stress development [50].

Total Flavonoid content (TFC) in methanolic extract was showed maximum quantity in fresh samples of *T. longipetalus* subsp. *longipetalus* (666.1±49μg/ml sample) shown S. Fig 2b. Fruit samples (fresh) highest value of flavonoid and ascorbic acid was detected in *T. terrestris* (566.1 ±5.1 μg/ml sample and 456.5±9.5 μg/g f. wt. respectively). In dry aerial parts *T. longipetalus* subsp. *longipetalus* flavonoid content gives maximum amount of TFC (495.4±16.4 μg /mL sample) followed by *F. bruguieri* var. *laxa* (395.1±37 μg/mL sample). While in seeds *P. harmala* showed maximum TFC (418.8±16.7μg/mL sample) followed by *F. bruguieri* var. *bruguieri* (391.8±5.12 μg/mL sample) Flavonoids are considered to be effective free radical scavengers in fruits, vegetables, and medicinal plants. Highest ascorbic acid reported in *P. harmala* shown S. Fig 2c. Ascorbic acid involved in a number of physiological processes. such as POD and SOD [51].The highest flavonoids and ascorbic acid content in fruits of *T. terrestris* and aerial parts of *T. longipetalus* subsp. *longipetalus* in this study validated its traditional medicine use and may be responsible to cure various ailments. *F. olivieri* shown S. Fig 2c gives the highest amount of tannins.

**S. Fig. 2:** Comparison of a) Total Phenolic Contents b) Total Flavonoid c) Ascorbic Acid Content. d) Tannin.

Total soluble protein and reducing sugar were high in fresh aerial parts of *P. harmala* shown in S Fig. 3a and b respectively. Earlier [52] isolated antioxidant protein from *P. harmala*. Seeds possessed antioxidant activity, and this activity was due to the presence of hydrophobic amino acids. *P. harmala* is one of the most frequently used medicinal plants to treat hypertension and cardiac disease worldwide [53]. The maximum total soluble proteins (168.3±6.3 mg/g dry wt.) were depicted in dry aerial parts of *Z. simplex*. No significant variation was observed in seed samples, the highest value of total soluble proteins (248.6±30 mg/g s. wt.) was found in *F. bruguieri* var. *rechingeri.* Total oxidant status (TOS) was lower in *F. indica* (1275±475 μM/g. f. wt.) shown in S Fig. 3d.

**S. Fig. 3:** Comparison of a) Total soluble proteins b) Reducing Sugar c) Total Oxidant Status d) Total Antioxidant Capacity.

Total flavonoid content was calculated in and expressed as μg/mL in methanolic extract using quercetin as standard (Table 2). Significant difference was observed among all selected species of Zygophyllaceae. In aerial parts *T. longipetalus* subsp. *longipetalus* flavonoid content gives maximum amount of TFC (495.4±16.4 μg /mL sample) followed by *F. bruguieri* var. *laxa* (395.1±37 μg/mL sample). While in seeds *P. hermala* showed maximum TFC (418.8±16.7μg/mL sample) followed by *F. bruguieri* var. *bruguieri* (391.8±5.12 μg/mL sample) (Table 3). Total Phenolic Content (TPC) was estimated, in aerial parts of selected taxa no significant TPC variation was found among (Table 4). In general, highest TPC was depicted in *T. longipetalus* subsp. *pterophorus* (63025±1725 μM/g. dry wt.) followed by *F. bruguieri* var. *rechingeri* (54600±1350 μM/g. dry wt.). Seeds of selected plant samples showed significant variation. Highest TPC was detected in *Z. propinquum* (69225±775μM/g. s. wt.) followed by *F. bruguieri* var. *rechingeri* (66850±3900 μM/g. s. wt.) shown in Table 3. No significant Tannin variation was detected among aerial parts as well as in seeds of all tested taxa (Table 2). However, highest amount of tannin was estimated in *T. longipetalus* subsp. *pterophorus* (40375±4125 μM/g dry wt.) followed by *Z. fabago* (39175±4825 μM/g. dry wt.). In seeds highest amount of tannin was estimated in *P. hermala* (47525±2575 μM/g s. wt.) followed by *Z. propinquum* (42625±175 μM/g. s. wt.) shown in Table 3. Ascorbic Acid was observed, in aerial parts of all selected taxa and significant difference found among studied taxa (Table 4). Highest value of AsA was found in *F. olivieri* (744.2±2.7 μg/g dry wt.) followed by *F. bruguieri* var. *bruguieri*. AsA content in seeds showed no significant variation. In general the highest AsA content was found in *F. bruguieri* var. *laxa* (740.8±2.19 μg/g s. wt.) given in Table 3. Alpha amylase activity in dry aerial parts of selected plants was assessed (Table 2) no significant variation was found among the taxa of Zygophyllaceae. However, highest α-amylase activity was found in *Z. simplex* (164.9 ±3.39 mg/g. dry wt.) followed by *Z. fabago* (153 ± 6.6 mg/g. dry wt.). In seeds of selected taxa significant variation was found among various taxa is shown in (Fig 19 f). The highest value was observed in *F. bruguieri* var. *laxa* (159.5±11.8 mg/g. s. wt.) followed by *F. ovalifolia* subsp. *pakistanica* (133.9±0.37 mg/g. s. wt.) given in Table 3. Significant variation of total soluble protein was observed among all tested taxa dry samples (Table 2). The maximum total soluble proteins (168.3±6.3 mg/g dry wt.) were depicted in *Z. simplex*. No significant variation was observed in seed samples, the highest value of total soluble proteins (248.6±30 mg/g s. wt.) was found in *F. bruguieri* var. *rechingeri* (Table 3). TAC was measured in aerial parts of selected taxa of Zygophyllaceae (Table 2), No significant variation was detected among all selected plants. In general maximum TAC was detected in *Z. eurypterum* (15.8±2.2 μM/g. dry wt.) followed by *Z. simplex* (15.7±2.33 μM/g.dry wt.). In seeds of selected taxa no significant variation was detected the maximum TAC was in *F. olivieri* (15.6±2.4 μM/g. s. wt.) second highest was in *F. bruguieri* var. *rechingeri* (15.5±2.5 μM/g. s. wt.) given in Table 3. Reducing Sugar was measured in aerial parts of all selected plants (Table 2) there was a significant difference found among all taxa. Highest value of reducing sugar was recorded in *Z. fabago* (7.47±0.2 mg/g. s. wt.) followed by *Z. propinquum* and with minimum difference (7.19 ±0.55 mg/g. s. wt.). In seeds highest value of reducing sugar was found in *T. terrestris* (7.9±0.1 mg/g. s. wt.) given in Table 3.

**Table 2: Phytochemical analysis in dry aerial parts of selected taxa of Zygophyllaceae**

**Table 3: Phytochemical analysis in seeds of selected taxa of Zygophyllaceae**

Lycopene content in dry aerial parts of various taxa was measured (Table 4). The higher value was found in *T. longipetalus* subsp. *pterophorus* (9.25± 1.8 mg/g dry wt.) followed by *T. terrestris* (8.12±0.01 mg/g dry wt.). Lycopene content in seeds of various taxa of Zygophyllaceae was investigated (Table 5). The higher value of lycopene was found in *F. ovalifolia* subsp. *pakistanica* (5.87±0.75 mg/g s. wt.) followed by *F. bruguieri* var. *rechingeri* (4.92±0.19 mg/g s. wt.). Chlorophyll a content was estimated in dry aerial parts of dry samples of various taxa of Zygophyllaceae (Table 4). The Chlorophyll a content ranged from maximum in *T. longipetalus* subsp. *pterophorus* (572.1±0.05 ug/g dry wt.) followed by *T. terrestris* (548.8±0.2 ug/g dry wt.) to minimum with significant difference in *Z. eurypterum* (92.5±8.0 ug/g dry wt.). Chlorophyll a content in seeds of various taxa (Table 5) ranged from maximum in *F. bruguieri* var. *rechingeri* (197±0.76 ug/g s. wt.) followed by *F. ovalifolia* subsp. *pakistanica* (171.2±5.3 ug/g s. wt.). Chlorophyll b content was assessed in dry aerial parts of different taxa of Zygophyllaceae (Table 4) Chlorophyll b content varied among various taxa. The Chlorophyll b content ranged from maximum in *T. longipetalus* subsp. *pterophorus* (228 ±63 ug/g dry wt.) followed by *T. terrestris* (170.2±1.6 ug/g dry wt.). In seeds of different taxa Chlorophyll b content varied ranged from maximum in *F. ovalifolia* subsp. *pakistanica* (70.7±11.2 ug/g s. wt.) next highest was *F. bruguieri* var. *rechingeri* (38±4.6 ug/g s. wt.) to minimum in *Z. propinquum* (2.9±0.01ug/g s. wt.) with a significant difference (Table 5). Total carotenoids were investigated in dry aerial parts of dry samples of different species of Zygophyllaceae (Table 4). Significantly highest value of total carotenoids was found in *T. longipetalus* subsp. *pterophorus* (38.6±0.6 mg/g dry wt.) followed by *T. terrestris* (38.2±0.02 mg/g dry wt.). In seeds samples significantly highest value of total carotenoids was found in *F. bruguieri* var. *rechingeri* (19.3±0.00 mg/g s. wt.) (Table 5) followed by *F. ovalifolia* subsp *pakistanica* (16.6±1.5 mg/g s. wt.). Total chlorophyll content of dried aerial parts was measured among different taxa of zygophyllaceae is shown in (Table 4). Maximum Content of total chlorophyll was depicted in *T. longipetalus* subsp. *pterophorus* (800±62.9 ug/g dry wt.) with significant difference. It was followed by *T. terrestris* (719±1.8 ug/g dry wt.). In seeds measured among different taxa of Zygophyllaceae is given in (Table 5). Significant variation was observed among various taxa. Maximum Content of total chlorophyll was depicted in *F. ovalifolia* subsp. *pakistanica* (242±16.6 ug/g s. wt.) Second highest in *F. bruguieri* var. *rechingeri* (235.3±3.9 ug/g s. wt.). The liquid chlorophyll supplements are important in enhancing energy, detoxification of liver and stomach and colon, eliminating the body and mouth odor, helping in anemia and aiding in the elimination of mould from the body. The carotenoids are vital because of their strong colours, antioxidant activity as well as their role as precursors of vitamin A. These can be used as safe chemicals for neutraceutical purposes and food supplementation [54].

**Table 4: Pigment analysis in dry aerial parts of selected taxa of Zygophyllaceae**

**Table 5: Pigment analysis in seeds of selected taxa of Zygophyllaceae**

Pigment analysis in fresh samples shown in S. Fig 4 a, b, c, d, and e gives the significant variation among the selected taxa. The chlorophyll contents (a, b and total), were higher in aerial parts of *T. longipetalus* subsp. *pterophorus* followed by *T. terrestris*. While carotenoids and lycopene were depicted maximum in *T. terrestris* followed by *T. longipetalus* subsp. *pterophorus.* The main carotenoids such as zeaxanthin, b-carotene, canthaxanthin, astaxanthin as well as lycopene are prepared synthetically in nutraceutical industry [55]. Furthermore, Lycopene is suggested as one of the efficient carotenoids group for quenching ability. The plants used in current study detected pigments in Zygophyllaceae taxa can be characterized for various above mentioned applications can be utilized as supplements or medicines.

**S. Fig.4:** Comparison of Pigment a) Lycopene content b) Chlorophyll a content c) Chlorophyll b content d) Total carotenoids e) Total chlorophyll content.

This study was carried out to assess the anti-inflammatory and anti-diabetic potential of naturally occurring medicinal plants of Zygophyllaceae from desert and semi-desert areas of Balochistan, Pakistan. The protein denaturation assay was followed for in *vitro* anti-inflammatory activity while anti-diabetic activity was determined by α-amylase inhibitory assay for 13 extracts of aerial parts and 12 seeds extracts. Selected plants also used in traditional medicines by the folks in Balochistan. Selection of plants was based on Ethnobotanical appraisal of local communities, their uses in folk medicines to cure various ailments like fever, cough, inflammation of organs, gonorrhea, urinary tract infection (UTI), diabetes and cancer etc. Previously secondary metabolites such as flavonoids and alkaloids derivatives were evaluated in Zygophyllaceae [56,57].

Inflammation in medical terms is demarcated as a pathophysiological procedure described by soreness, fever, swelling of body parts, loss of function and discomfort [58]. The inhibitory effect among different plants of Zygophyllaceae on albumin denaturation is shown in (Fig. 5). Significant inhibition of albumin was observed among different taxa. *Z. eurypterum* seeds depicted maximum inhibition with the highest value (96.85±1.85 % Inh.). Seeds of *T. longipetalus* subsp. *pterophorous* revealed the next highest value (95.85±2.85 %), Seeds of *T. terrestris* exhibited the (95.35±3.35% Inh.) activity. *T. longipetalus* subsp*. longipetalus* depicted (91.1±1.1% of Inh.). Earlier in the fruit of *T. terrestris* significant anti-inflammatory activity was evaluated [59]. All species of *Fagonia* also exhibited anti-inflammatory activity significant and comparable with the standard drug diclofenac sodium. In seeds extract highest value were found in *F. bruguieri* var. *laxa* (88.9 ± 4.9% Inh). *F. ovalifolia* subsp. *pakistanica* showed (84.8±0.8%) of inhibition. *F. olivieri* and *F. bruguieri* var*. rechingeri* showed (80.9 ±0.9 and 79.7±0.7% Inh. respectively). Seeds of *P. harmala* constituted (76.4±1.4 % Inh.) activity for inhibition. Seeds of *F. bruguieri* var. bruguieri (66.2±1.2% Inh.) revealed less inhibition when compared with the standard drug. Previously anti-inflammatory activity of *F. cretica* was studied by [60]. Minimum inhibition of albumin was observed in seeds of *Z. propinquum* (15.79±0.20% Inh.) in comparison with all studied taxa. Previously, *Nitraria schoberi* from Zygophyllaceae fruit extract has been found to have anti-inflammatory effects [61] that are quite in line with presently observed activities for seeds and aerial parts. Aerial parts of all the studied taxa of Zygophyllaceae also exhibited significant inhibition of albumin. *T. longipetalus* subsp*. pterophorus* revealed maximum inhibition (90.1±2.1 % Inh.) followed by *T. terrestris* and *Fagonia bruguieri* var*. laxa* (89.9±0.9 and 89.4±0.9 % Inh. respectively) anti-inflammatory inhibition. *Z. fabago* and *F. olivieri* indicated (88.3 ± 1.3 % Inh.) inhibition in both plants. *F. ovalifolia* subsp. *pakistanica* (endemic to the region) revealed (87.1±1.1% Inh.) albumin inhibition activity. *T. longipetalus* subsp. *longipetalus, Z. simplex* and *F. bruguieri* var*. bruguieri* exhibited inhibition activity as (86.5 ± 0.5, 83.2 ±1.2 and 80.4 ±1.2 % Inh. respectively). *P. harmala* and *Z. propinquum* aerial parts also showed significant inhibition as compared with standard drug (79.6±0.6 and 75.9±1.9 % Inh.). *F. indica* (49.6±0.3 % Inh.) depicted less inhibition when compared with all other taxa as well as standard drug. Previously chloroform and methanol extract of aerial parts of *T. terrestris* exhibited significant anti-inflammatory activities at a dose of 200 mg/kg [59]. All examined *Fagonia* spp. in present study exhibited therapeutic potential especially *F. bruguieri* var. *laxa* suggested to be used as anti-inflammatory as well as anti-diabetic agent authenticating its folk use. Earlier, *F. cretica* was identified for its high therapeutic effects against various types of hematological, liver disorders, neurological and inflammatory conditions. Furthermore, aqueous extract is one of the seventeen ingredients in Norm acid syrup used in the treatment of high acidity and gastritis [62]. Furthermore, the endemic taxa of the region i.e. *F. ovalifolia* subsp*. pakistanica* also exhibited high anti-inflammatory potential. Previously *F. longipina* traditionally used as a preventive for cancer, also used for the treatment of inflammation of the urinary tract [48]. *F. schweinfurthii* plant extract gel could be developed as a therapeutic agent for wound healing and anti-inflammatory properties [63].

**Fig 5:**
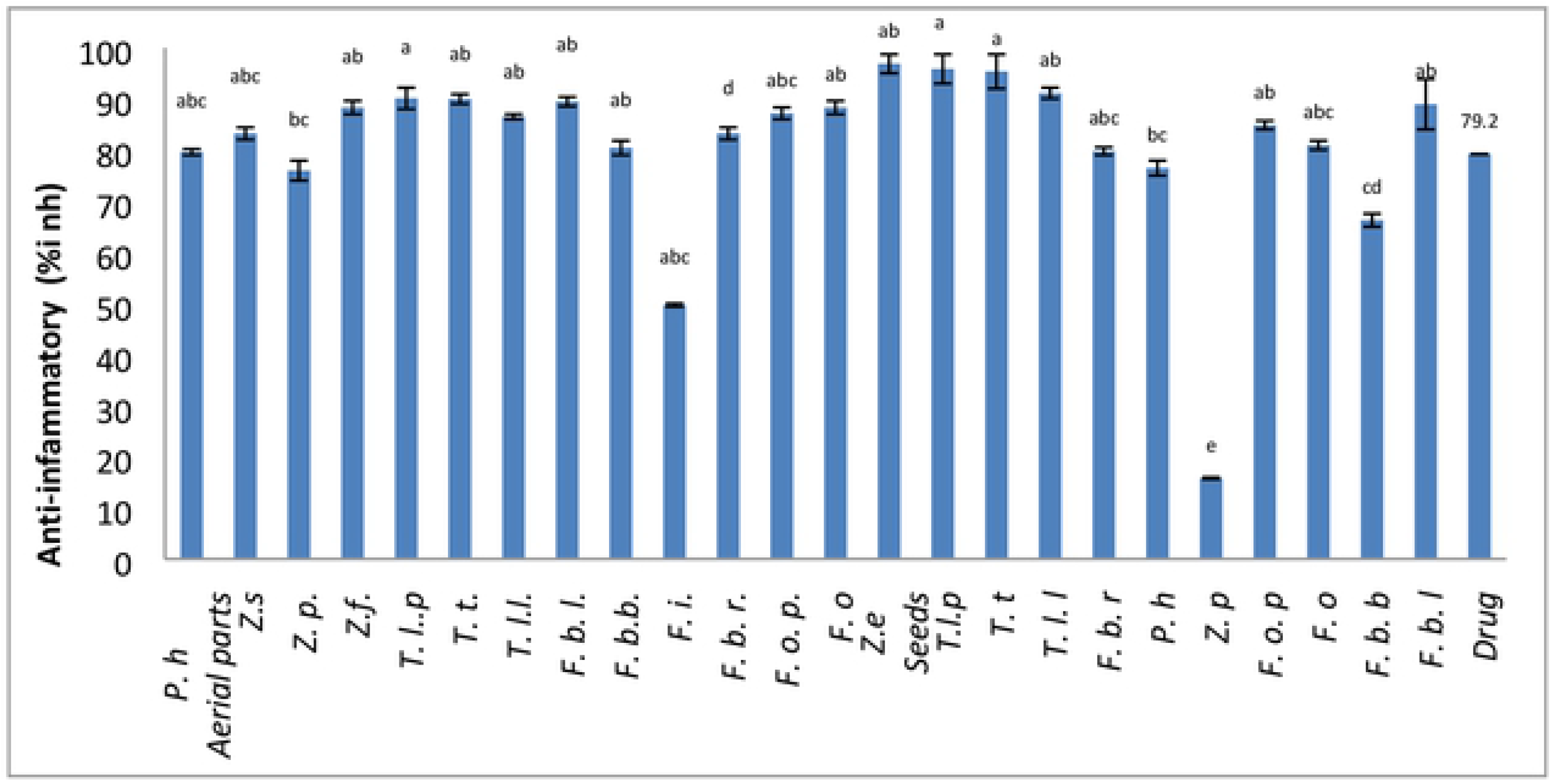
Comparison of Anti-inflammatory activity among different taxa of Zygophyllaceae. Data are presented as mean values ± SEM (n = 3). Statistical analysis: ANOVA test and Tukey (HSD). The different letters above the values in the same column indicate significant differences with Tolerance: 0.0001.

Anti-diabetic activity was analyzed by using amylase inhibition assay. Seeds and aerial parts of selected species revealed significant differences in anti-diabetic activity when compared with standard drug Metformin (Fig.6). The seeds of *T. longipetalus* subsp. *longipetalus* (85.65±0.34% Inh.) exhibited the highest anti-diabetic activity among all selected species. It was followed by seeds of *Z. eurypterum* (83.63±0.63% Inh.), next highest value was found in *T. terrestris* (82.8±0.1%). Seeds of *Z. propinquum* indicated (81.5±0.4%) activity. *F. bruguieri* var. *rechingeri* depicted (80.9±1.9 %) anti-dibetic activity. *F. bruguieri* var *bruguieri* revealed (80.3±0.6 %) enzyme inhibition in the seeds of *F. bruguieri* var. *laxa* (79.5±1.0%) inhibition was found. Seeds of *T. longipetalous* subsp. *pterophorus* reported (77.7±0.2%) enzyme inhibition activity. The endemic plant of the region *F. ovalifolia* subsp*. pakistanica* depicted (76.5±0.4%) anti-diabetic activity. *P. harmala* showed (71.4±1.4%) anti-diabetic activity. Seeds of *F. olivieri* exhibited (69.8±0.8%) enzymatic inhibitory activity Seeds of *P. harmala* used as anti-inflammatory, antidiabetic [64]. Same therapeutic potential of seeds of *P. harmala* was attained in the current study.

**Fig 6:**
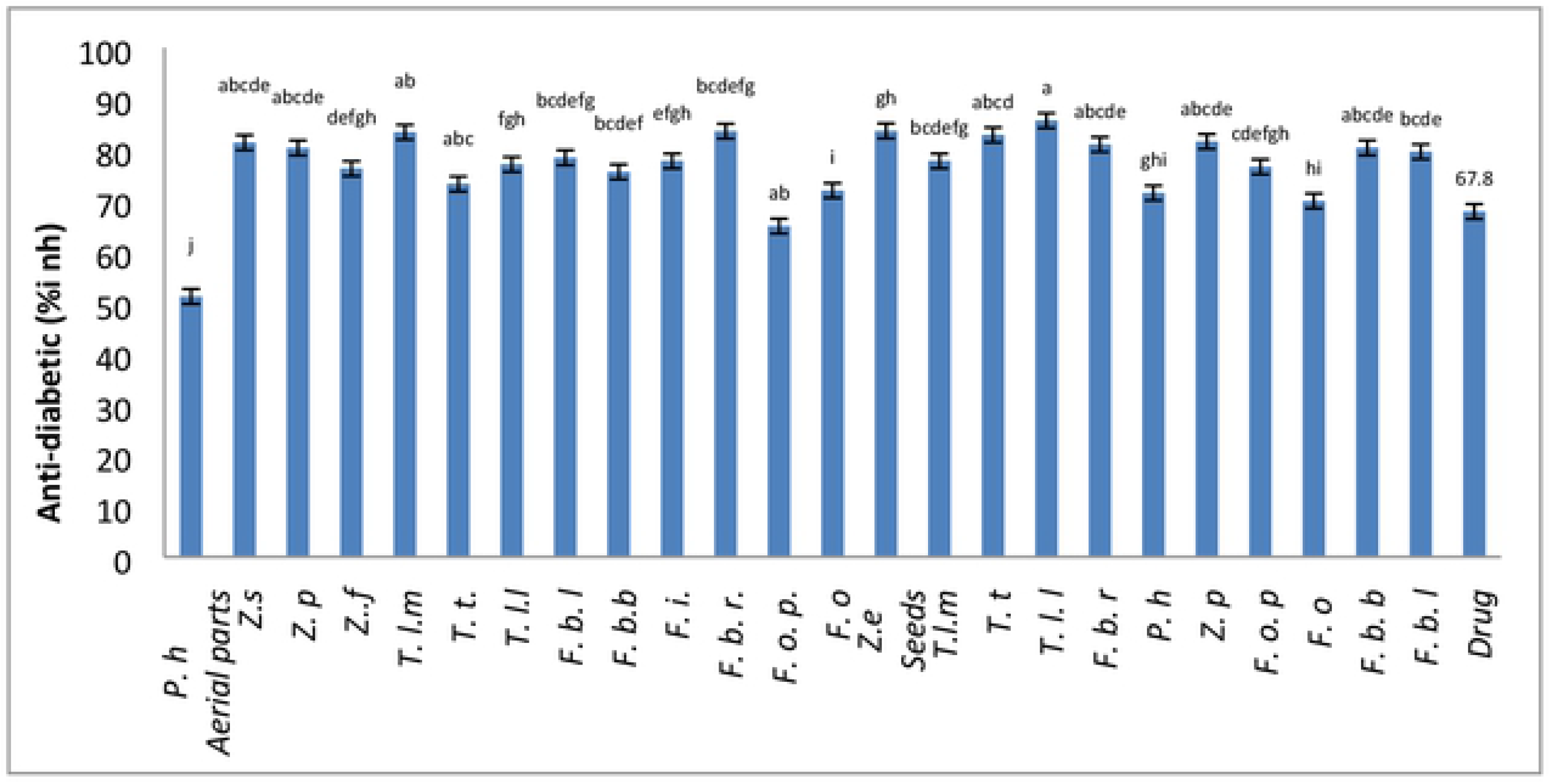
Comparison of Anti-diabetic activity of different taxa of Zygophyllaceae Data are presented as mean values ± standard deviation (n = 3). Statistical analysis: ANOVA test and Tukey (HSD). The different letters above the values in the same column indicate significant differences with Tolerance: 0.0001

Aerial parts of selected taxa also exhibited significant enzyme inhibition activity when compared with standard drug. Highest value was found in *F. bruguieri* var. *rechingeri* (83.59±0.40 % Inh.). Second highest value was in *T. longipetalus* subsp*. pterophorus* (83.37±0.62 % Inh.). Next highest value was found in *Z. simplex* (81.3±1.3% Inh.) followed by *Z. propinquum* (80.2±1.2 % Inh.) succulent plants. *F. bruguieri* var *laxa* depicted (78.3±1.1 % Inh.) anti-diabetic activity. *F. indica* showed (77.7±0.2 % Inh.) activities followed by *T. longipetalus* subsp. *longipetlus* (77 ±1.06% Inh.) inhibition. Aerial parts of *Z. fabago* shows (76*.2*±1.*2*% Inh.) inhibitory effect. *F. bruguieri* var. *bruguieri* showed (75.5±0.5 % Inh.) enzymatic inhibitory activity. Aerial parts of endemic plant *F. ovalifolia* subsp*. pakistanica* also showed significant inhibition of amylase enzyme (64.9±0.05% Inh.) Earlier reported *F. indica* alone or combined with *Aloe vera* can be used as a natural blood glucose lowering agent [65]. In present research the other examined *Fagonia* species are potentially active therapeutic agents in comparison with *F. indica*. *P. harmala* and *Z. gaetulum* frequently used to treat hypertension and diabetes mellitus [53].

## Conclusion

Phytochemical screening of all selected of Zygophyllaceae revealed that these plants have significant potential of enzymatic, non-enzymatic activities and other phytochemicals like flavonoids, total phenolic compounds, tannins, and pigments. All the taxa proved to have natural antioxidants and could not only be used to treat various ailments but also contribute in the prevention of degenerative diseases and manufacturing of new drugs. Research findings could be utilized for isolation of potential phytopharmacological active compounds from these wild medicinal plants for future research. Species such as *Z. eurypterum, T. longipetalus* subsp*. longipetalus, T. terresetris*, *F. bruguieri* and *F. ovalifolia* subsp. *pakistanica* had greater potential for identification, isolation and purification of novel therapeutic agents.

